# Virulence and genomic diversity among clinical isolates of ST1 (BI/NAP1/027) *Clostridioides difficile*

**DOI:** 10.1101/2023.01.12.523823

**Authors:** Qiwen Dong, Huaiying Lin, Marie-Maude Allen, Julian R. Garneau, Jonathan K. Sia, Rita C. Smith, Fidel Haro, Tracy McMillen, Rosemary L. Pope, Carolyn Metcalfe, Victoria Burgo, Che Woodson, Nicholas Dylla, Claire Kohout, Anitha Sundararajan, Evan S Snitkin, Vincent B. Young, Louis-Charles Fortier, Mini Kamboj, Eric G. Pamer

## Abstract

*Clostridioides difficile (C. difficile)*, a leading cause of nosocomial infection, produces toxins that damage the colonic epithelium and results in colitis that varies from mild to fulminant. Variation in disease severity is poorly understood and has been attributed to host factors (age, immune competence and intestinal microbiome composition) and/or virulence differences between *C. difficile* strains, with some, such as the epidemic BI/NAP1/027 (MLST1) strain, being associated with greater virulence. We tested 23 MLST1(ST1) *C. difficile* clinical isolates for virulence in antibiotic-treated C57BL/6 mice. All isolates encoded a complete Tcd pathogenicity locus and achieved similar colonization densities in mice. Disease severity varied, however, with 5 isolates causing lethal infections, 16 isolates causing a range of moderate infections and 2 isolates resulting in no detectable disease. The avirulent ST1 isolates did not cause disease in highly susceptible Myd88^-/-^ or germ-free mice. Genomic analysis of the avirulent isolates revealed a 69 base-pair deletion in the N-terminus of the *cdtR* gene, which encodes a response regulator for binary toxin (CDT) expression. Genetic deletion of the 69 base-pair *cdtR* sequence in the highly virulent ST1 R20291 *C. difficile* strain rendered it avirulent and reduced toxin gene transcription in cecal contents. Our study demonstrates that a natural deletion within *cdtR* attenuates virulence in the epidemic ST1 *C. difficile* strain without reducing colonization and persistence in the gut. Distinguishing strains on the basis of *cdtR* may enhance the specificity of diagnostic tests for *C. difficile* colitis.

## Introduction

*Clostridioides difficile* is a Gram-positive, spore-forming anaerobic bacterium, and the leading cause of nosocomial infections in the United States.^1–3^ Infections are acquired by oral ingestion of *C. difficile* spores, which are prevalent in the environment and can survive for extended periods of time on contaminated surfaces. Upon ingestion, *C. difficile* spores germinate, produce toxins and cause colitis and, in severe cases, can result in mortality. The major virulence factors of *C. difficile* are toxins A (*tcdA*) and B (*tcdB*), which are encoded in the Pathogenicity Locus (PaLoc).^4,5^ These toxins glycosylate and thereby inactivate host GTPases, triggering the death of intestinal epithelial cells and lead to gut inflammation.^6^

*C. difficile* species is comprised of hundreds of strain types across more than 6 phylogenetic clades. PCR- and sequencing-based approaches, including PCR-ribotyping (RT) and multilocus sequencing typing (MLST/ST), have been used to characterize *C. difficile* strain types. More recently, whole-genome sequencing (WGS) has greatly contributed to our understanding of *C. difficile* diversity, evolution, and epidemiology.^7^ Almost two decades ago, the BI/NAP1/027 strain, characterized as ST1 by MLST, emerged as a cause of severe nosocomial outbreaks and increased *C. difficile* infection (CDI) incidence in North America and Europe. Since then, the prevalence of ST1 has declined but it remains among the most frequently isolated strains in hospital and community-acquired CDI cases in the US.^2,8–11^ The ST1 *C. difficile* strain encodes the additional CDT toxin (encoded by *cdtA* and *cdtB* and also referred to as binary toxin), which is an ADP-ribosyltransferase that modifies actin and disrupts cellular cytoskeleton organization.^12^ The ST1 strains have higher MICs to several antibiotics, most notably fluroquinolones, and produce higher amounts of toxins A and B compared to non-ST1 strains.^13,14^ The relative virulence of the ST1 strain is controversial, however, with some studies demonstrating clinical disease severities similar to other strains.^15–17^ Host factors can impact the severity of CDI, including underlying diseases, immune competence and microbiome composition.^18,19^ Whether genetic variants of ST1 explain diverse disease manifestations is unknown.

To determine intra-strain type virulence diversity, we used an antibiotic-treated mouse model of *C. difficile* infection to test a panel of PaLoc- and CdtLoc-encoding ST1 *C. difficile* clinical isolates to quantify disease severity.^20^ Clinical *C. difficile* isolates with identical PaLoc caused a range of disease severities, with two isolates causing no detectable disease in antibiotic-treated wild type, germ-free mice or Myd88-defecient mice. We identified a 69-bp deletion in the *cdtR* gene of these two avirulent isolates, which encodes a LytTR family response regulator that regulates CDT expression. The 69-bp deletion in the *cdtR* leads to reduced CDT toxin and PaLoc gene expression, resulting in loss of virulence and confirming previous studies implicating CdtR as regulator of CDT and Tcd toxins expression.^21^ Our study is the first to describe virulence diversity within a single strain type and demonstrates the critical role of CdtR for ST1 *C. difficile* virulence.

## Results

### Clinical ST1 *C. difficile* isolates demonstrate variable severities in mice

We focus on a group of 23 *C. difficile* isolates belonging to ribotype 027 epidemic strains (here referred to as ST1) isolated from patients with diarrhea during a molecular surveillance program at Memorial Sloan Kettering Cancer Center 2013-2017.^22^ Wholegenome Illumina sequencing of these isolates allows us to compare them to the public collections. We plotted a UMAP analysis of the presence or absence of unique coding sequences (annotated proteins or un-annotated protein clusters) across top 10 STs of *C. difficile* strains in Patric (date: Feb. 10 2021).^23^ Different STs cluster individually and our ST1 isolates overlap with other ST1 *C. difficile* included in the analysis, confirming their strain type **(Figure 1A)**. These 23 isolates demonstrate high genome-wide similarity by average nucleotide identity (ANI) score above 99.8% and encode identical Pathogenicity Locus (PaLoc) sequences **(Figure S1A-S1B)**. To study if close-related *C. difficile* isolates may have variable virulence, mice treated with antibiotics (metronidazole, neomycin, vancomycin in drinking water with clindamycin intraperitoneal injection) were orally infected with each of these isolates at a dose of 200 spores and *C. difficile* pathogenicity was monitored throughout a 7-day-timecourse **(Figure 1B)**. Mice infected with different ST1 isolates displayed a spectrum of disease severity, including variable weight loss and mortality **(Figure 1C and Figure S1C)**. The widely used ST1 lab strain R20291 was included in parallel for virulence comparison. Within our ST1 collection, 5 isolates resulted in mortality in mice. A few isolates caused more severe weight loss than R20291, including ST1-49, ST1-11 and ST1-12, while most ST1 isolates caused moderate and non-lethal infections. Two isolates, ST1-75 and ST1-35 demonstrated no impact on mouse body weights. No apparent colonization deficiency was observed in any of these isolates **(Figure S1D)**. The variable pathogenicity induced by a group of ST1 isolates with identical PaLoc suggested additional regulatory mechanisms of *C. difficile* virulence. Therefore, we sought to examine other genomic factors that are responsible for attenuated virulence of *C. difficile* isolates ST1-75 and ST1-35.

**Figure 1.**
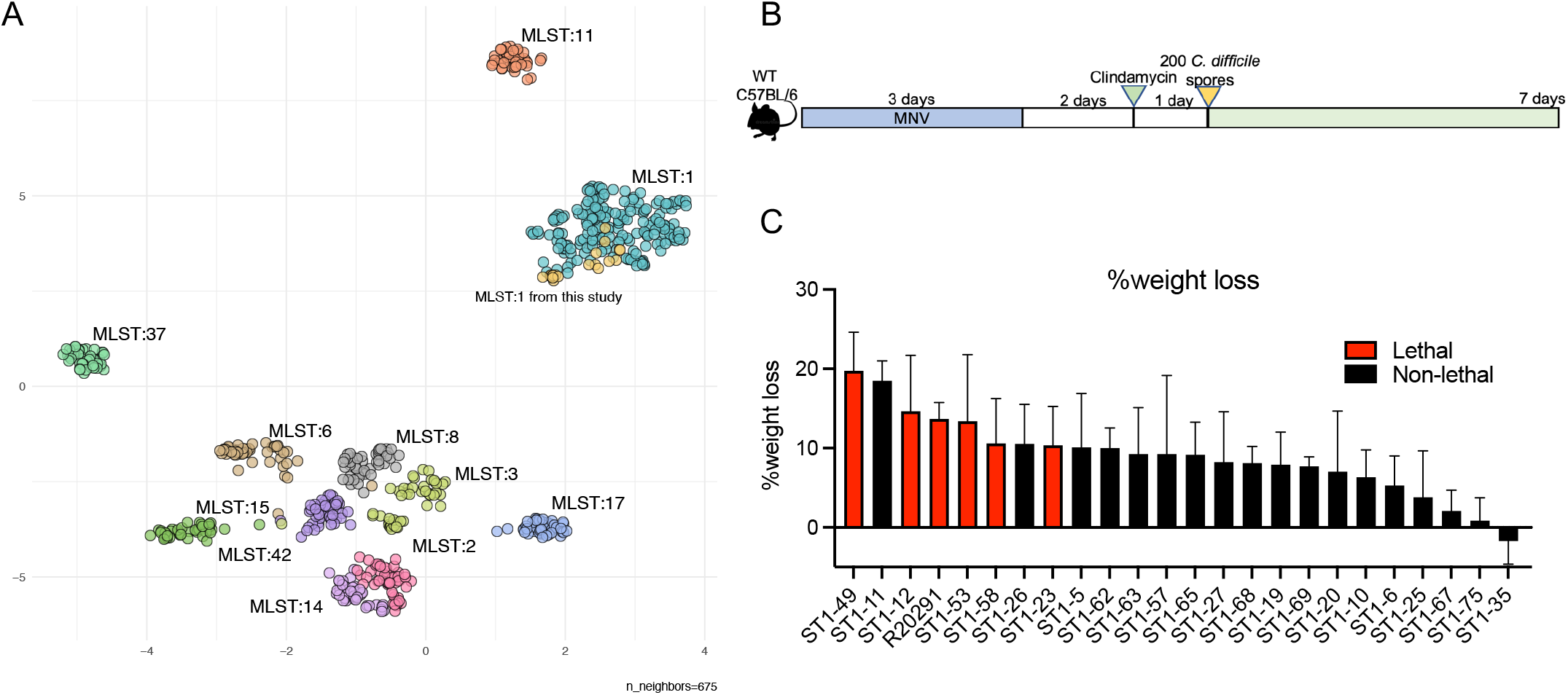
Clinical ST1 *C. difficile* isolates demonstrated variable virulence in mice treated with antibiotics. (A) Plot of the a UMAP analysis of the presence or absence of unique coding sequences (annotated proteins or un-annotated protein clusters) across top 10 STs of *C. difficile* strains in Patric. (B) Mouse experiment schematic: wildtype C57BL/6 mice were treated with metronidazole, vancomycin, and neomycin (MNV, 0.25 g/L for each) in drinking water for 3 days and followed by one intraperitoneal injection of clindamycin (200 μg/mouse) 2 days after antibiotic recess. Then, mice were inoculated with 200 *C. difficile* spores via oral gavage. Daily body weight and acute disease scores were monitored for 7 days post infection. (C) %Max weight loss to baseline were calculated using the lowest weights within 7 days post infection divided by day 0 weights. N (Number of mice per strain-infected group) = 5-8 except for ST1-62 and ST1-68, which have 2 mice per group.

### Two ST1 *C. difficile* isolates demonstrate avirulent phenotype

Among the *C. difficile* isolates that we examined using antibiotic-treated mice, two isolates, ST1-75 and ST1-35, caught our attention due to their strikingly attenuated phenotypes **(Figure 1B)**. Almost no weight loss was observed throughout the 7-day-timecourse, and low acute disease scores were displayed in mice infected with ST1-75 or ST1-35, in comparison to mice infected with R20291 **(Figure 2A-2B and 2D-2E)**. This avirulent phenotype was not due to colonization deficiency of ST1-75 or ST1-35, as the fecal CFU recovered from the mice infected with these two isolates were comparable to R20291-infected mice on both early and late days post-infection **(Figure 2C and 2F)**. Fecal levels of TcdA and TcdB were also measured, and similar levels were seen in the feces at day +1 post-infection from mice infected with ST1-75 and ST1-35, compared to R20291**(Figure 2G-2H)**.

**Figure 2.**
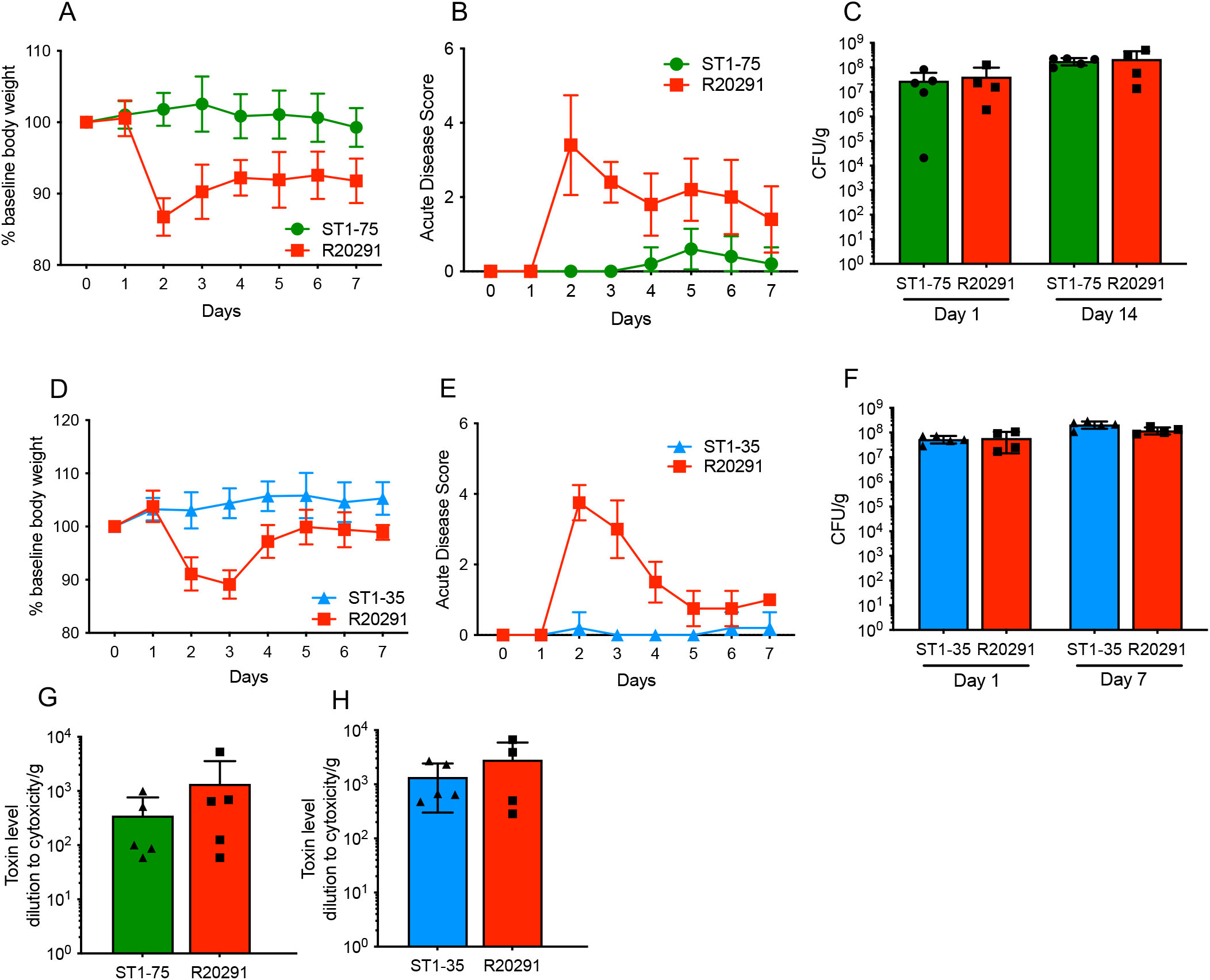
Two isolated clinical strains of *C. difficile* have no virulence in mice treated with antibiotics. Wildtype C57BL/6 mice (n=3-5 per group) were treated with metronidazole, vancomycin, and neomycin (MNV, 0.25 g/L for each) in drinking water for 3 days and followed by one intraperitoneal injection of clindamycin (200 μg /mouse) 2 days after antibiotic recess. Then, mice were inoculated with 200 *C. difficile* spores via oral gavage. Daily body weight and acute disease scores were monitored for 7 days post infection. (A, D) %Weight loss to baseline of mice infected with indicated strains. (B, E) Acute disease scores comprising weight loss, body temperature drop, diarrhea, morbidity of mice infected with indicated strains. (C, F) Fecal colony-forming units measured by plating on selective agar on indicated days.(G-H) Fecal toxins measured by CHO cell rounding assay 1 day post infection.

To further investigate this avirulent phenotype, we inoculated ST1-75 into MyD88^-/-^ mice, which lack the adaptor protein for Toll-like receptor signaling.^24^ MyD88^-/-^ mice fail to recruit neutrophils to the colonic tissue during early stages of *C. difficile* infection, and display markedly increased susceptibility to *C. difficile* induced colitis.^25^ Here, MyD88^-/-^ mice were treated with antibiotics and infected with either ST1-75 or R20291. Mice infected with R20291 quickly succumbed to infection 2 days after spore inoculation, whereas all MyD88^-/-^ mice infected with ST1-75 survived the experiment with minimal weight loss or disease scores **(Figure 3A-3B)**. Consistent with our results with wildtype mice, no deficiencies of colonization or toxin production were observe dday+1 post infection of MyD88^-/-^ mice **(Figure 3C-3D)**. These data suggest that the attenuation of the avirulent strain is independent of MyD88-mediated host innate immunity.

**Figure 3.**
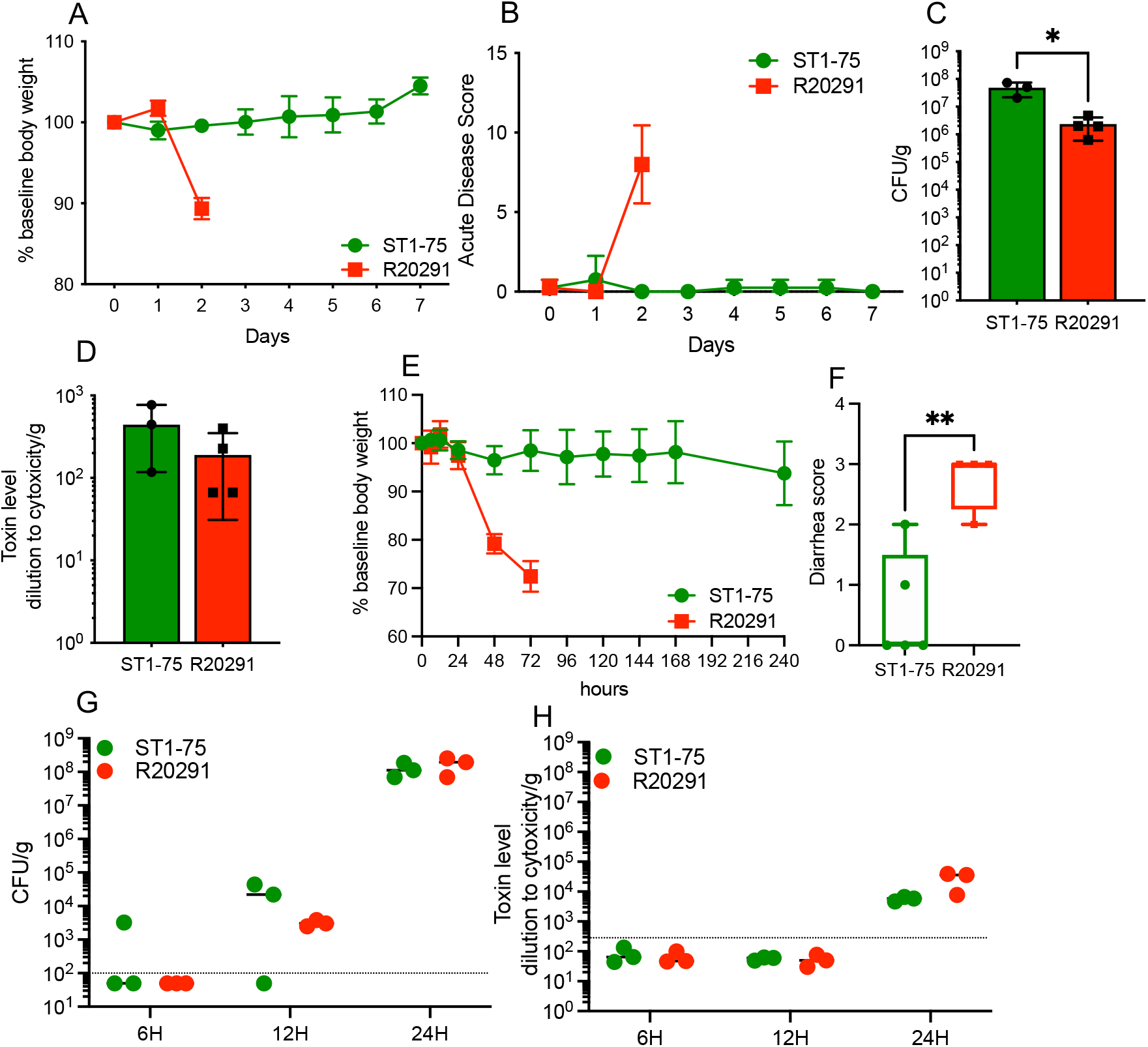
Avirulent *C. difficile* strain demonstrates no virulence in innate immune deficient mice and germ-free mice. (A-D) MyD88^-/-^mice (n=4 per group) were treated with MNV and clindamycin before orally administered with 200 spores of *C. difficile* strains. Daily body weight and acute disease scores were monitored for 7 days post infection. (A) %Weight loss to baseline of mice infected with indicated strains. (B) Acute disease scores comprising weight loss, body temperature drop, diarrhea, morbidity of mice infected with indicated strains. (C) Fecal colony-forming units measured by plating on selective agar 1 day post infection. (D) Fecal toxins measured by CHO cell rounding assay 1 day post infection. (E-H) Germ-free mice (n=3 to 5) orally administered with 200 spores of indicated *C. difficile* strains. Daily body weight and acute disease scores were monitored for 10 days post infection. (E) %Weight loss to baseline of mice infected with indicated strains. (F) Diarrhea scores of mice infected with indicated strains 2 days post infection. (G) Fecal colony-forming units measured by plating on selective agar at 6, 12, and 24 hours post infection. (H) Fecal toxins measured by CHO cell rounding assay at 6, 12, and 24 hours post infection. Statistical significance was calculated by Unpaired t-test, * p < 0.05, ** p < 0.01.

Germ-free mice are highly susceptible to *C. difficile* infection because they microbiome-mediated colonization resistance against *C. difficile*.^26,27^ To investigate whether the gut microbiome renders ST1-75 avirulent, germ-free mice were infected with ST1-75 or R20291. Similarly, we observed no mortality or weight loss in the mice with ST1-75 infection, whereas mice infected with R20291 quickly lost weight and died (or >20% weight loss) **(Figure 3E)**. Milder diarrhea was observed in mice with ST1-75 compared to R20291 **(Figure 3F).** We observed no differences in colonization or fecal toxins between ST1-75 and R20291 up to 24 hours post infection **(Figure 3G-3H)**. Similar attenuation was also seen in ST1-35 infected germ-free mice **(Figure S2)**. In contrast, isolates that demonstrated relatively mild pathogenicity in antibiotic-treated mice, such as ST1-25 and ST1-67 **(Figure 1B)**, led to severe weight loss and diarrhea in germ-free mice **(Figure S2),** reaffirming the protective role of the gut microbiome during *C. difficile* infection. However, the attenuation of ST1-75 and ST1-35 in mice is independent of the gut microbiome.

### Novel prophages identified in avirulent strains do not impact ST1 *C. difficile* virulence

We next sought to determine the genetic factors that may abrogate *C. difficile* virulence in ST1-75 and ST1-35. Fully circularized genomes of 14 ST1 isolates were successfully obtained using Nanopore and Illumina hybrid assembly, and pangenomic analysis was conducted on these 14 genomes and R20291 using Anvi’o pangenomics workflow.^28^ A group of gene clusters that is unique to ST1-75 and ST1-35 stood out, which are enriched for phage-related genes **(Figure 4A)**. We then applied PHASTER, a tool for phage identification in bacterial genomes, to discover two unique prophages in the genomes of ST1-75 and ST1-35. One prophage resides on a 41-kb plasmid in ST1-75 and ST1-35 with 4-5 copies per cell and here is named as phiCD75-2 **(Figure S3A)**. Blasting phiCD75-2 found high similarities to reported *C. difficile* phages phiCD38-2 (99.8% identity) and phiCDHS1 (94.7% identity).^29–32^ In addition, a unique 54-kb segment was found as inserted into the chromosomal DNA of ST1-75 and ST1-35 around position 1.29 Mbp and here is named as phiCD75-3 **(Figure S3A)**. PhiCD75-3 does not show high similarity to any described *C. difficile* phages to date.

**Figure 4.**
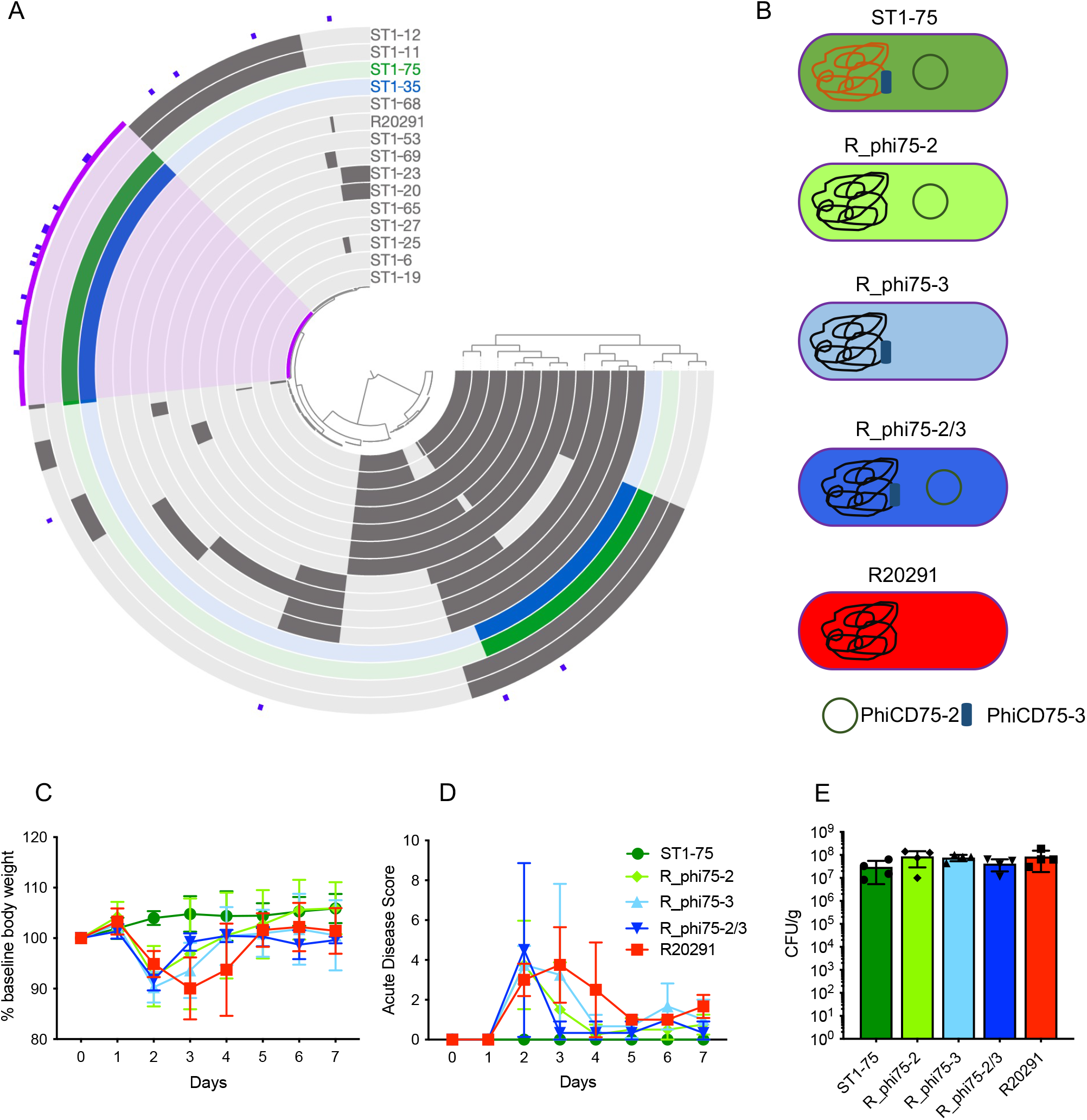
Prophages identified in avirulent *C. difficile* mildly impact virulence in mice treated with antibiotics. (A) Anvi’o plot displaying accessory genomes of ST1 isolates. Highlighted gene clusters in purple are unique to ST1-35 and ST1-75. Blue dashes (outmost layer) indicate phage-related genes by NCBI COG. (B) Schematic of mutant strains made using R20291 *C. difficile* strain. (C-E) Wildtype C57BL/6 mice (n=4 per group) were treated with MNV and clindamycin as preciously described. Then, mice were inoculated with 200 *C. difficile* spores via oral gavage. Daily body weight and acute disease scores were monitored for 7 days post infection. (C) %Weight loss to baseline of mice infected with indicated strains. (D) Acute disease scores comprising weight loss, body temperature drop, diarrhea, morbidity of mice infected with indicated strains. (E) Fecal colony-forming units measured by plating on selective agar 1 day post infection.

Lysogenic bacteriophages have been identified in many *C. difficile* genomes, and play an important role in shaping *C. difficile* evolution. However, their roles in *C. difficile* biology, especially virulence, are not well-characterized.^33,34^ To investigate the potential role of these two prophages on *C. difficile* virulence, we induced lytic phage particles of phiCD75-2 and phiCD75-3 from ST1-75 culture and infected R20291 to generate R20291 lysogens harboring these prophages. We were able to generate R20291 derivatives carrying phiCD75-2, phiCD75-3 or both prophages in their genomes **(Figure 4B)**. Whole-genome sequencing of lysogenic R20291 strains confirmed that phiCD75-3 was inserted *in situ* as in ST1-75 at 1.29 Mbp. Antibiotic-treated mice infected with R20291 lysogenic strains **(Figure 4B)** followed the curve of virulent infection as 10% weight loss and 4-6 disease scores were seen during the peak of symptomatic infection **(Figure 4C-4D).** The seemingly faster recovery in the lysogens was not a reproducible finding. Similar levels of colonization and toxin production were also observed **(Figure 4E and S3B-S3D)**. Here, we discover two prophages in avirulent strains ST1-75 and ST1-35, that are not present in R20291 or other ST1 strains from our collection of. However, these two prophages do not appear to impact the virulence of R20291 in antibiotic-treated mice.

### Mutations in the *cdtR* gene eliminate ST1 *C. difficile* virulence in mice

Lysogenic R20291 strains with either or both prophages did not recapitulate the avirulent phenotype of ST1-75 or ST1-35. A closer look at the chromosomal genomes of ST1-75 and ST1-35 led us discover a common mutation in their *cdtR* gene, which was reported previously as a transcriptional regulator for binary toxin (CDT) genes, *cdtA* and *cdtB*.^35^ A unique 69-bp deletion was found in the *cdtR* gene of ST1-75 and ST1-35, leading to an inframe deletion of 23 amino acids **(Figure 5A)**. To investigate if there is a possible loss of function of CdtR resulted from the deletion, we accessed the transcriptional level of *cdtB* in mouse cecum following infection of ST1-75 or R20291. More than a 2-log reduction of *cdtB* transcripts was observed in ST1-75 group **(Figure 5B)**, suggesting an important role of these 69 base pairs for a fully functional *cdtR* gene. Next, we applied CRISPR-mediated genome editing approach to generate CdtR mutants using parental R20291 strain to study the contribution of CdtR to *C. difficile* virulence **(Figure S4A and 5A)**. In accordance with a previous report^21^, knocking out *cdtR* either by deleting the whole gene (CdtRKO8.1 and CdtRKO10.3), or introducing a proximal premature stop codon (CdtRstop4.2 and CdtRstop8) led to a loss of pathogenicity in antibiotic-treated mice **(Figure S4B-S4C)**, confirming a critical role of CdtR for *C. difficile* virulence. Moreover, deleting the exact same 69-bp region, as in ST1-75/35, in the *cdtR* of R20291 (CdtRmut6.1 and CdtRmut8.1) again eliminated the virulence of *C. difficile* **(Figure 5C-5D)**. Thus, loss of the 69-bp in *cdtR* explains the avirulence phenotype of ST1-75/35. On the other hand, colonization of these CdtR mutants, assessed by CFU at day+1 post infection, was comparable to that of R20291 **(Figure S4D and S4F)**, suggesting that CdtR is not required for colonization. Interestingly, while the fecal levels of TcdA and TcdB of CdtR mutants were comparable to R20291 in the early phase (day+1 post-infection), a significantly reduced level at a later time point (7-days post infection) was observed upon infection of CdtR mutants **(Figure S4E and 5E)**, supporting a role of CdtR in regulating PaLoc toxins production. Further, infecting germ-free mice with CdtRmut6.1 results in no weight loss or diarrhea, perfectly recapitulating ST1-75 and ST1-35 phenotypes in germ-free mice **(Figure 5F-5G and Figure S4G)**. Collectively, CRISPR-edited CdtR mutant strains mimic phenotypes of ST1-75 and ST1-35 in mice, demonstrating that *cdtR* gene is necessary for *in vivo* virulence. Furthermore, the 69-bp region in *cdtR*, which is deleted in ST1-75/35, is necessary for proper CdtR function through mechanisms yet to be determined.

**Figure 5.**
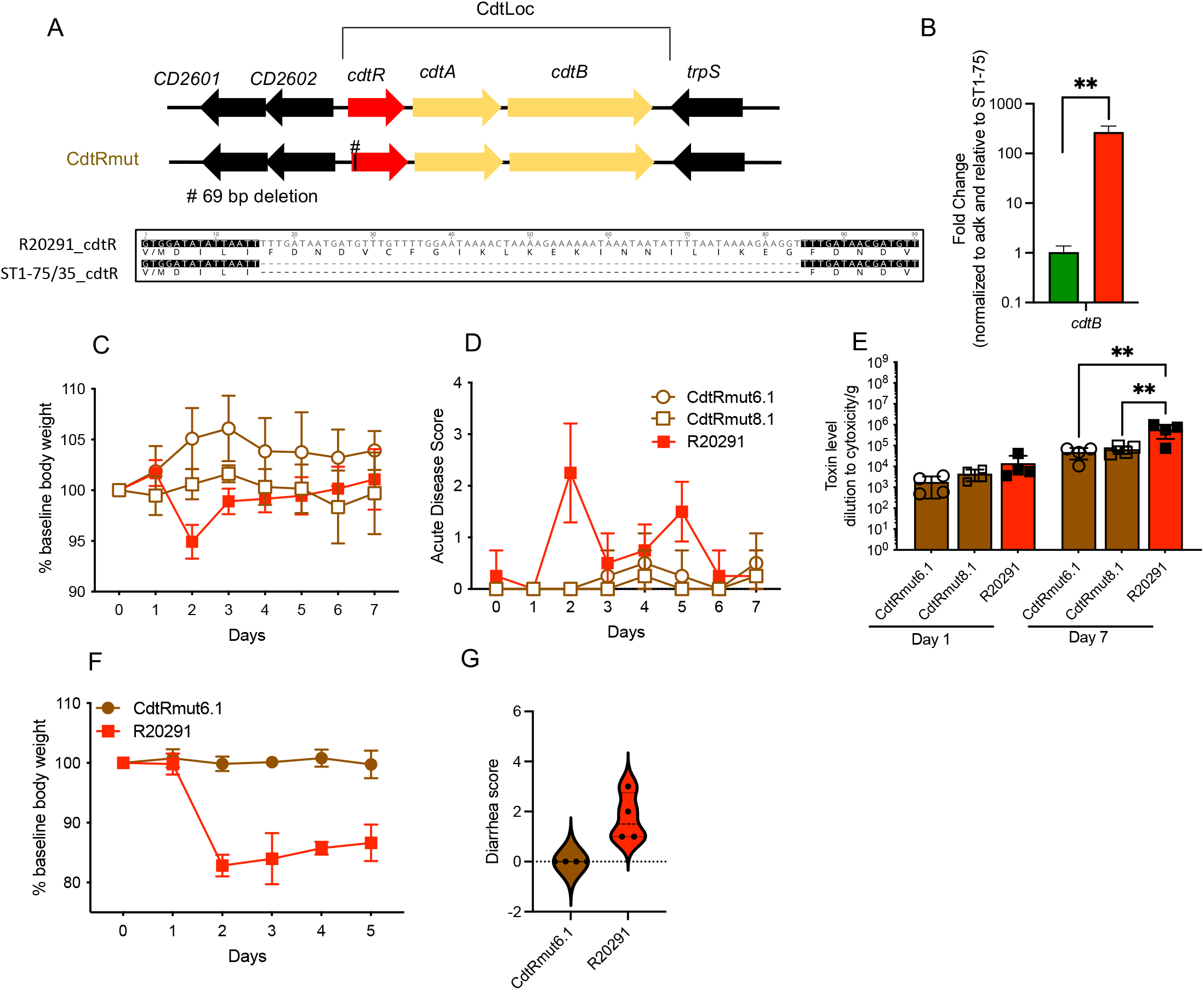
Binary toxin regulator *cdtR* contributes to *C. difficile* virulence in mice. (A) Deletion identified in ST1-35/75 and schematic of *cdtR* mutants generated using R20291 *C. difficile* strain. (B) Germ-free mice (n=3 per group) orally administered with 200 spores of indicated *C. difficile* strains. Binary toxin gene *cdtB* transcripts were measured by RT-qPCR on cecal contents harvested at 24 hours post infection. Transcripts were normalized to the *adk* and fold change is relative to ST1-75 condition. (C-E) Wildtype C57BL/6 mice (n=3 to 5 per group) were treated with MNV and clindamycin as preciously described. Then, mice were inoculated with 200 *C. difficile* spores via oral gavage. Daily body weight and acute disease scores were monitored for 7 days post infection. (C) %Weight loss to baseline of mice infected with indicated strains. (D) Acute disease scores comprising weight loss, body temperature drop, diarrhea, morbidity of mice infected with indicated strains. (E) Fecal toxins measured by CHO cell rounding assay on indicated days. (F-G) Germ-free mice (n=4) orally administered with 200 spores of indicated *C. difficile* strains. Daily body weight and were monitored for 5 days post infection. (F) %Weight loss to baseline of mice infected with indicated strains. (G) Diarrhea scores of mice infected with indicated strains 3 days post infection. Statistical significance was calculated by One-way ANOVA, * p <0.05, ** < p< 0.01.

### Mutations in *cdtR* reduce PaLoc toxin transcription *in vivo*

CdtR mutants produce significantly reduced fecal toxins at a later time point (7-days post infection) **(Figure 5E and S4E)**, a phenotype that was confirmed in ST1-75 and ST1-35 **(Figure S5A-S5B)**. To examine whether the 69-bp deletion in *cdtR* impacts PaLoc toxin production, we harvested cecal contents from germ-free mice infected with ST1-75 or R20291. In contrast to fecal toxin levels, we observed a significantly reduced cecal toxin level in mice infected with ST1-75 compared to R20291 **(Figure 6A)**. This was not due to a slightly lower CFU of cecal ST1-75 in germ-free mice **(Figure S5C-S5D)**. We further validated the reduced toxin production in cecal content by RT-qPCR and we observed a 50-fold reduction of the *tcdA* and *tcdB* transcripts in the cecum of mice infected with ST1-75 **(Figure 6B)**. Additionally, transcriptions of other PaLoc genes including *tcdE* and *tcdR* were also reduced in the cecum of mice infected with ST1-75 **(Figure 6B)**. *TcdE* is a putative holin that mediates toxin secretion.^36–38^ Reduced *tcdE* likely further impacts the amount of toxins that may reach to the intestinal epithelium. TcdR is a positive regulator of the PaLoc^39,40^ and is likely the common target of CdtR, which results in the observed downregulation of many PaLoc genes. Interestingly, *cdtR* transcripts were comparable between ST1-75 and R20291, suggesting that the 69-bp deletion does not impact the transcripts stability but leads to a nonfunctioning product. These results were further confirmed with CdtRmut6.1 strain, though to a lesser extent **(Figure S5E)**. Collectively, we demonstrated that a natural mutation found in *cdtR* of two ST1 clinical isolates results in reduced binary toxin production, reduced PaLoc toxin production (and likely secretion) in cecum of infected mice, and attenuated *C. difficile* virulence, independent of host innate immunity, colonization burden, microbiome constitution, or any noticeable impact of incidentally discovered prophages within these strains. This difference of toxin production in cecum at 24-hour post infection is however, not reflected in feces in parallel, but could be reflected at a later time point, likely due to cumulative differences over time.

**Figure 6.**
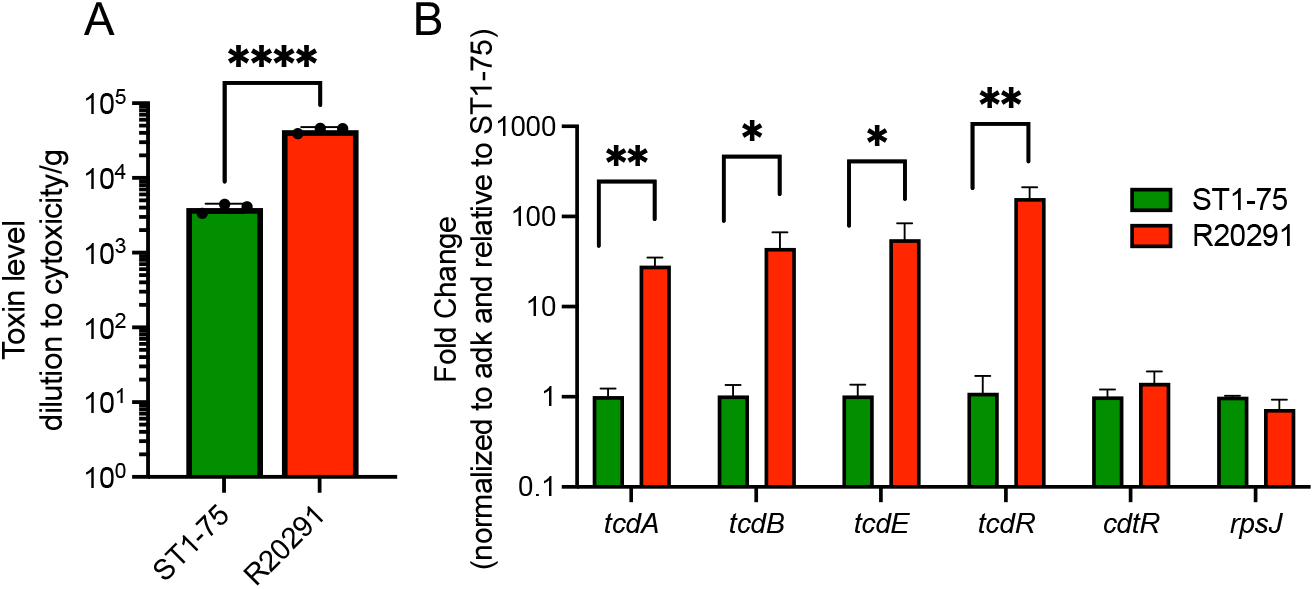
CdtR regulates PaLoc toxins transcription *in vivo*. Germ-free mice (n=4) were orally administered with 200 spores of indicated *C. difficile* strains and cecal contents were harvested at 24 hours post infection. (A) Cecal toxins measured by CHO cell rounding assay. (B) Indicated gene transcripts were measured by RT-qPCR. Transcripts were all normalized to the *adk* and fold change is relative to ST1-75 for each of the genes. Statistical significance was calculated by Unpaired t-test, * p < 0.05, ** p < 0.01, **** p < 0.0001.

### *cdtR* is versatile and more prevalent than *cdtA* and *cdtB*

Our data support a regulatory role of CdtR outside CdtLoc, so we hypothesize that CdtR may evolve to impact virulence beyond regulating CDT binary toxins. To test this possibility, we surveyed the presence of *cdtR*, *cdtA* and *cdtB* in two major *C. difficile* clinical collections.^23,41^ As expected, the majority of clade 2 strains, including the epidemic ST1/RT027 strains, contains the CdtLoc with the presence of all three genes. Other subgroups of *C. difficile* strains, including MLST5 and MLST11 were also reported to encode CDT **(Figure 7A)**.^42,43^ Unexpectedly, many strain types of *C. difficile* that were reported as CDT-negative also encode *cdtR*, such as MLST2, MLST8 from clade 1 **(Figure 7A)**. The higher prevalence of *cdtR* over *cdtA* and *cdtB* supports the possibility that CdtR functions beyond regulating CDT. Additional work is needed to evaluate the functions of CdtR in these CDT-negative strains. On the contrary, the presence of *cdtR* in CDT-positive strains may lose its function, such as in MLST11 strains, where a premature stop codon was found and results in a *cdtR* pseudogene^42^, as well as here in the case of ST1-75/35. To evaluate the prevalence of *cdtR* mutations that may lead to a loss of function, we aligned all *cdtR* genes in MLST1 strains from the two described collections and found a few strains having similar truncations at the proximal end and may have lost CdtR function, yet we did not find the exact same deletion as in ST1-75/35 within almost 500 strains **(Figure S6A)**.

**Figure 7.**
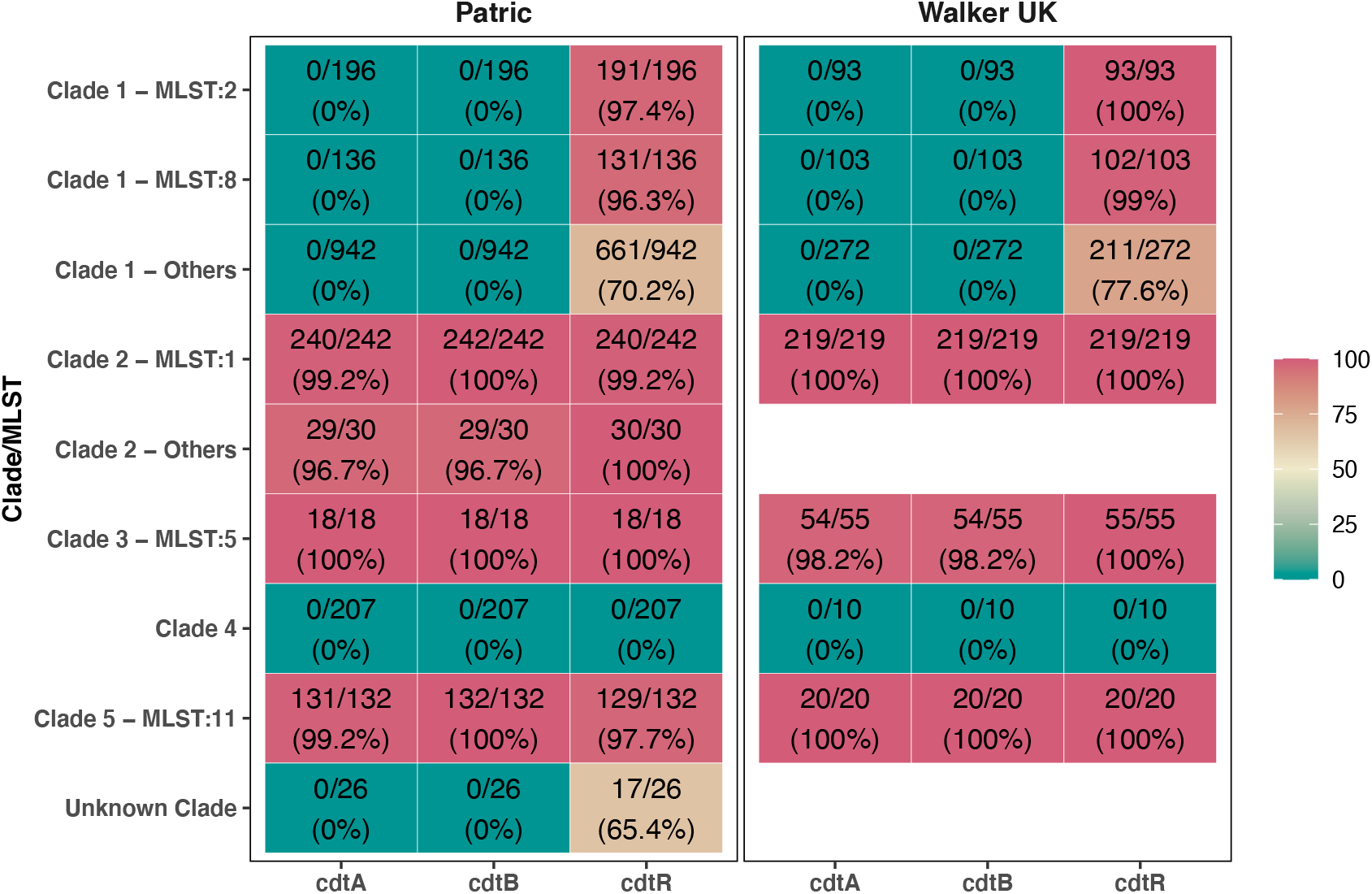
Binary toxin regulator *cdtR* is prevalent in clinical *C. difficile* isolates. Binary toxin *cdtA, cdtB* and *cdtR* from R20291 were used as query to BLAST against the assembled contigs. Hits with at least 85% identity and 85% coverage of the query were considered a valid match. Numbers of match in total and percentages are presented.

The high similarity between ST1-75 and ST-35 genetically and phenotypically led us reason whether they are clonal. We performed a core-genome SNP analysis across all ST1 isolates from our collection with R20291 as the reference. ST1-75 and ST1-35 shared all SNPs when compared to the genome of R20291 **(Figure S6B).** Additional clinical evidence supports ST1-35 and ST1-75 being clonal strains isolated from two patients that shared the same hospital room a few days apart **(Figure S6C)**. Our survey on CdtLoc genes suggest that this locus is very versatile during evolution and *cdtR* is more prevalent than *cdtA* and *cdtB*, which may regulate virulence beyond CDT.

## Discussion

Mouse models are valuable tools to study how *C. difficile* strain variations may result in variable disease severities, thanks to the advantages of their identical genetic, immune background and controlled microbiome compositions. Here we focused on a group of clinical *C. difficile* isolates belonging to the RT027/MLST1, with high genomic similarity, that all encode PaLoc and CdtLoc. We found that these similar *C. difficile* isolates caused variable disease severities in mice and that a very specific mutation in the *cdtR* gene rendered two clinical isolates, ST1-75 and ST1-35, avirulent. Avirulence was solely dependent on the *cdtR* mutation, as we obtained similar observations using MyD88^-/-^ mice, and germ-free mice, which was also further validated with CRISPR-edited *cdtR* mutants. Lower transcripts of binary toxin gene *cdtB*, toxins A *tcdA*, toxin B *tcdB*, together with other PaLoc genes including regulator *tcdR* and putative holin *tcdE*, were observed in mouse cecum infected with the CdtR mutants. Our data support a critical role of CdtR in regulating *C. difficile* toxin production and secretion, which is essential to ST1 virulence. However, all the other ST1 isolates in this study encoded an intact CdtLoc with wildtype *cdtR*, whose variations in virulence are likely attributable to alternative mechanisms.

The presence of a binary toxin locus has been associated with epidemic strains and hypervirulence of *C. difficile*.^44,45^ CDT belongs to the family of ADP-ribosylating toxins that consist of two components: CDTa (*cdtA*), the enzymatic active ADP-ribosyltransferase which modifies cellular actin, and CDTb (*cdtB*), the binding component facilitates CDTa translocation. However, despite knowing their enzymatic activities, experimental evidence is very limited to support critical roles of CDT in *C. difficile* virulence.^46^ CDTb was reported to induce TLR2-dependent pathogenic inflammation, which suppresses a protective eosinophilic response and enhances virulence of RT027 strains, however, *C. difficile* lacking CDTb still causes acute disease in mice.^47^ On the other hand, CdtR, as the transcriptional regulator of *cdtA* and *cdtB*^6,35,48^, has been previously linked to PaLoc toxin production^21^, suggesting a role as a major virulence regulator. Here, we demonstrated a critical role of CdtR as a determinant of *C. difficile* virulence within ST1 strains. A natural 69-bp deletion in *cdtR* that was found in two clinical isolates can reverse the virulence of a wildtype strain, by downregulating the expression of PaLoc genes and binary toxin genes. Additionally, higher prevalence of *cdtR* over *cdtA* or *cdtB* was found while surveying CdtLoc on clinical isolates from public databases. This suggests CdtR may have evolved to function beyond regulating *cdtA* and *cdtB*. Systematically examining the target genes of CdtR may give us insights on its additional functions, which may also help unveil the mechanisms by which CdtR regulates the PaLoc genes.

ST1-75 and ST1-35 are avirulent in susceptible mouse models despite producing toxins, albeit at reduced levels. This is intriguing because it is well appreciated that toxin expression is necessary for *C. difficile* virulence.^46,49^ However, our data indicate that toxin production is not sufficient for causing CDI. The amount of toxin being produced and released is likely impact the development of disease. The patients from whom we isolated ST1-75 or ST1-35 had an overall mild clinical assessment, whose symptoms may be attributable to causes other than *C. difficile* infection. Current CDI diagnoses largely depend on the detection of *TcdB* gene or toxin B positivity in feces and may lead to overdiagnosis of CDI. We, together with other reports, suggest the importance of quantifying toxins to evaluate CDI cases.^50–52^ Incorporating adjunctive biomarkers, such as IL-1β, better distinguishes CDI from asymptomatic carriage and non-CDI diarrhea.^53^ Here, CdtR regulates both toxin production and secretion, and is essential for *C. difficile* virulence in mice, suggesting it may serve as an adjunctive biomarker for CDI diagnosis.

Apart from characterizing CdtR, we also identified two prophages in ST1-75 and ST1-35. Prophages have been identified in many *C. difficile* genomes, and play important roles in shaping *C. difficile* evolution.^33^ While prophages are highly prevalent in *C. difficile*, little is known about how prophages impact *C. difficile* biology. A couple of pioneering studies have shown that prophages can affect *C. difficile* gene expression, impacting toxin production.^29,54,55^ In this study we identified two prophages in ST1-75/35 and named them phiCD75-2 and phiCD75-3. By making R20291 lysogenic strains harboring either or both prophages, we observed minimal impacts on *C. difficile* virulence by neither of the prophages in antibiotic-treated mice. PhiCD38-2 was shown to increase PaLoc gene expression and toxin production in some RT027 isolates, but not in all of them, suggesting that the genetic background influences the impact of a newly acquired prophage.^29^ This may explain why phiCD75-2 (a phiCD38-2 derivative) did not increase toxin production ST1-75. Certain phages also impact phase variation of the cell surface protein, biofilm formation, and carry genes involved in quorum sensing, inferring their roles in bacterial fitness.^30,56,57^ It would be very intriguing to investigate how phiCD75-2 and phiCD75-3 may impact *C. difficile* fitness, including gene expression, antibiotic resistance, and interspecies competition.

In summary, we demonstrate that ST1 *C. difficile* clinical isolates with identical PaLoc display variable virulence *in vivo*. Among them, two clonal clinical isolates, ST1-75 and ST1-35, were avirulent in mice, due to a 69-bp deletion mutation in their *cdtR* genes. These data suggest that specific *cdtR* genetic variants within the same strain type may predict disease occurrence and severity. Routine detection of these variants may enhance the specificity of NAATs for CDI diagnosis. Our data also corroborate recent clinical observations that toxin detection is unreliable as the sole criterion to distinguish between *C. difficile* infection and colonization.

## Experimental model and subject details

### *C. difficile* clinical isolate collection and classification

Toxigenic *C. difficile*-positive stool specimens were collected at Memorial Sloan Kettering Cancer Center between 2013-2017. *C. difficile* isolates were recovered by plating onto brain heart infusion (BHI) agar plates supplemented with yeast extract, L-cysteine (BHIS), and the antibiotics D-cycloserine and cefoxitin (BHI and yeast extract were from BD Biosciences, and the other components were from Sigma-Aldrich) in an anaerobic chamber (Coylabs). Individual colonies that were able to grow in the presence of these antibiotics and that had the characteristic phenotype of *C. difficile* were selected, isolated, and subjected to whole-genome sequencing and MLST classification.^58^

### Mouse husbandry

Wild-type C57BL/6 mice, aged 6 to 8 weeks, were purchased from the Jackson Laboratories. MyD88^-/-^ mice were maintained in augmentin (0.48 g/L and 0.07 mg/L of amoxicillin and clavulanate respectively) in the drinking water in specific-pathogen-free (SPF) facility at the University of Chicago. Germ-free C57Bl/6J mice were bred and maintained in plastic gnotobiotic isolators within the University of Chicago Gnotobiotic Core Facility and fed ad libitum autoclaved standard chow diet (LabDiets 5K67) before transferring to BSL2 room for infection. Mice housed in the BSL2 animal room are fed irradiated feed and provided with acidified water. All mouse experiments were performed in compliance with University of Chicago’s institutional guidelines and were approved by its Institutional Animal Care and Use Committee.

## Method details

### *C. difficile* spore preparation and numeration

*C. difficile* sporulation and preparation was processed as described previously^59^ with minor modifications. Briefly, single colonies of *C. difficile* isolates were inoculated in deoxygenated BHIS broth and incubated anaerobically for 40-50 days. *C. difficile* cells were harvested by centrifugation and five washes with ice-cold water. The cells were then suspended in 20% (w/v) HistoDenz (Sigma, St. Louis, MO) and layered onto a 50% (w/v) HistoDenz solution before centrifugating at 15,000 × g for 15 minutes to separate spores from vegetative cells. The purified spores pelleted at the bottom were then collected and washed for four times with ice-cold water to remove traces of HistoDenz, and finally resuspended in sterile water. Prepared spores were heated to 60°C for 20 min to kill vegetative cells, diluted and plated on both BHIS agar and BHIS agar containing 0.1% (w/v) taurocholic acid (BHIS-TA) for numeration. Spore stocks for mouse infection were verified to have less than 1 vegetative cell per 200 spores (as the infection dose).

### Virulence assessment of clinical isolates in mice

SPF mice were treated with antibiotic cocktail containing metronidazole, neomycin and vancomycin (MNV) in drinking water (0.25g/L for each antibiotic) for 3 days, 2 days after removing MNV, the mice were received one dose of clindamycin (200 μg/mouse) via intraperitoneal injection. Mice were then the next day infected with 200 *C. difficile* spores via oral gavage. Germ-free mice were infected with 200 *C. difficile* spores via oral gavage without antibiotic treatments.

Following infection, mice were monitored and scored for disease severity by four parameters^60^: weight loss (> 95% of initial weight = 0, 95%–90% initial weight = 1, 90%–80% initial weight = 2, < 80% = 3), surface body temperature (> 95% of initial temp= 0, 95%–90% initial temp = 1, 90%–85% initial temp = 2, < 85% = 3), diarrhea severity (formed pellets = 0, loose pellets = 1, liquid discharge = 2, no pellets/caked to fur = 3), morbidity (score of 1 for each symptoms with max score of 3; ruffled fur, hunched back, lethargy, ocular discharge).

### Quantification of fecal colony forming units

Fecal pellets or cecal content from *C. difficile* infected mice were harvested and resuspended in deoxygenated phosphate-buffed saline (PBS), diluted and plated on BHI agar supplemented with yeast extract, taurocholic acid, L-cysteine, D-cycloserine and cefoxitin (CC-BHIS-TA) at 37°C anaerobically for overnight.^61^

### Cell-based assay to quantify fecal and cecal toxin

The presence of *C. difficile* toxins was determined using a cell-based cytotoxicity assay as previously described with minor modifications.^61^ Briefly, Chinese hamster ovary cells (CHO/dhFr-, ATCC#CRL-9096) were incubated in a 96-well plate overnight at 37°C. Tenfold dilutions of supernatant from resuspended fecal or cecal content were added to CHO/dhFr-cells, incubated overnight at 37°C. Cell rounding and death was scored the next day. The presence of *C. difficile* toxins was confirmed by neutralization by antitoxin antisera (Techlab, Blacksburg, VA). The data are expressed as the log10 reciprocal value of the last dilution where cell rounding was observed.

### DNA extraction, RNA extraction and reverse transcription

Fecal DNA was extracted using DNeasy PowerSoil Pro Kit (Qiagen), and RNA was isolated from cecal contents using RNeasy PowerMicrobiome Kit (Qiagen) according to the manufacturer’s instructions, respectively. Complementary DNA was generated using the QuantiTect reverse transcriptase kit (Qiagen) according to the manufacturer’s instructions.

### Quantitative polymerase chain reaction (qPCR)

Quantitative PCR was performed on genomic DNA or complementary DNA using primers (listed in Table 1) with PowerTrack SYBR Green Master Mix (Thermo Fisher). Reactions were run on a QuantStudio 6 pro (Thermo Fisher). Relative abundance was normalized by ΔΔCt.

**Table 1.**
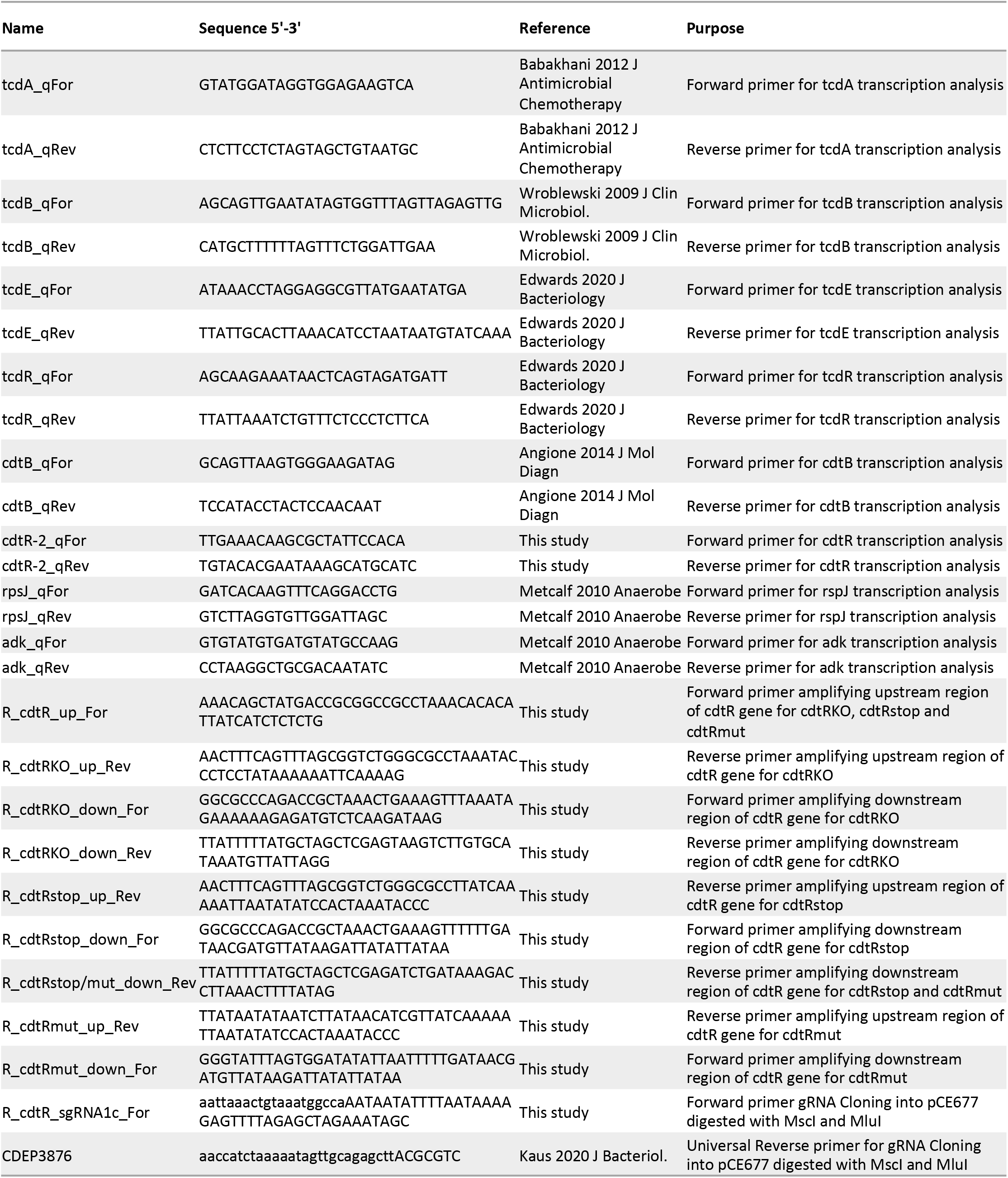
Oligonucleotides used in this study.

### Generation of *C. difficile cdtR* mutants using CRISPR

CRISPR editing on *C. difficile* strains R20291 was performed as described in.^62^ The primers were listed in Table 1^63,65,67,69,71^. Briefly, donor regions for homology were generated by separately amplifying regions ~500 bp upstream and ~500 bp downstream of the target of interest. The resulting regions were cloned into pCE677 between NotI and XhoI sites by Gibson Assembly. Geneious Prime (v11) was used to design sgRNAs targeting each deleted target. sgRNA fragments were then amplified by PCR from pCE677, using an upstream primer that introduces the altered guide and inserted at the MscI and MluI sites of the pCE677-derivative with the appropriate homology region. Regions of plasmids constructed using PCR were verified by Sanger sequencing. Plasmids were then passaged through NEBturbo *E. coli* strain before transformation into *Bacillus subtilis* strain BS49. The CRISPR-Cas9 deletion plasmids which harbor the oriT (Tn916) origin of transfer, were then introduced into *C. difficile* strains by conjugation.^64^ *C. difficile* colonies were then screened for proper mutations in the genomes by PCR and correct clones were further validated by whole-genome sequencing.

### Whole-genome sequencing and assembly

DNA was extracted using the QIAamp PowerFecal Pro DNA kit (Qiagen). Libraries were prepared using 100 ng of genomic DNA using the QIAseq FX DNA library kit (Qiagen). Briefly, DNA was fragmented enzymatically into smaller fragments and desired insert size was achieved by adjusting fragmentation conditions. Fragmented DNA was end repaired and ‘A’s’ were added to the 3’ends to stage inserts for ligation. During ligation step, Illumina compatible Unique Dual Index (UDI) adapters were added to the inserts and prepared library was PCR amplified. Amplified libraries were cleaned up, and QC was performed using Tapestation 4200 (Agilent Technologies). Libraries were sequenced on an Illumina NextSeq 500 or MiSeq platform to generate 2×150 or 2×250 bp reads respectively. Illumina reads were assembled into contigs using SPAdes^66^ and genes were called and annotated using Prokka (v1.14.6).^68^

Samples for Nanopore and Illumina hybrid assemblies were extracted using the NEB Monarch Genomic DNA Purification Kit. DNA was QC’ed using genomic Tapestation 4200. Nanopore libraries were prepared using the Ligation Sequencing Kit (SQK-LSK109), the Native Barcoding Expansions 1-12 (EXP-NBD104) and 13-24 (EXP-NBD114), and the NebNext Companion Module for Oxford Nanopore Technologies (E7180S). The shearing steps and first ethanol wash were eliminated to ensure high concentrations of long fragments. Using R9.4.1 flow cells, libraries were run on a MinION for 72 hours at −180V. The Nanopore and Illumina hybrid assemblies were completed using Unicycler (v0.4.8)^70^ either with the untrimmed or trimmed Illumina reads. The assemblies with less number of circularized contigs were used for genome analysis.

### Multiple sequence alignment for Pathogenicity loci

Illumina whole-genome of twenty-five *C. difficile* strains including two reference genome: R20291 (accession: FN545816.1) and CD630 (accession: NC_009089.1), along with 23 in-house strains ST1-10, ST1-11, ST1-12, ST1-19, ST1-20, ST1-23, ST1-25, ST1-26, ST1-27, ST1-35, ST1-49, ST1-5, ST1-53, ST1-57, ST1-58, ST1-6, ST1-62, ST1-63, ST1-65, ST1-67, ST1-68, ST1-69, and ST1-75 were included in the pathogenicity locus (PaLoc) analysis. The PaLoc region of R20291 (NCBI accessions NC_013316, 706,660 – 725,022 bp) were extracted as the query to BLAST^72^ against a local database of all the above genomes. Hits with at least 85% query coverage and 85% percent identity were extracted and multiple sequence alignment were performed using Geneious Prime 2022.0.1 with default settings to compare their nucleotide differences.

### Binary toxin genes prevalence analysis

*C. difficile* isolates (N=827) from BioProject PRJEB4556 were downloaded from NCBI, and assembled into contigs using SPAdes.^66^ A collection of 2143 *C. difficile* genomes from Patric (date: Feb. 10 2021)^23^ were also downloaded. MLST was determined on those contigs by mlst.^73^ ST type with less than 3 isolates were removed. Binary toxin *cdtA, cdtB* and *cdtR* from R20291 (NCBI accessions NC_013316) were used as query to BLAST^72^ against the assembled contigs, and hits with at least 85% identity and 85% coverage of the query are considered a valid match.

### UMAP (Uniform Manifold Approximation and Projection) analysis

A subset of isolate contigs of 199 ST1, 50 ST2, 50 ST3, 49 ST6, 50 ST8, 50 ST11, 42 ST14, 50 ST15, 50 ST17, 50 ST37 and 50 ST42, totaling 690 isolates were selected from the above Patric collection. They were all sequenced by short read technology, and they are the top 10 abundant ST groups except ST1 in the Patric collection. Genes were called and annotated from their contigs using Prokka (v1.14.6).^68^ By combining the 23 isolates from this study, we constructed a matrix of 731 isolates by 8025 annotated genes and hypothetical protein clusters. Specifically, hypothetical protein clusters were formed by clustering hypothetical proteins at 50% identity using cd-hit.^74,75^ Any protein sequences that were at least 50% similar fall into an artificially cluster. UMAP analysis was performed on the basis of the presence/absence of the genes/hypothetical protein clusters by setting the n_neighbors parameter to 675.

### Core-genomes SNPs analysis on ST1 isolates

SNP analysis was done on the 23 ST1 isolates from our collection against R20291 using snippy (v4.6.0).^76^ Then the core SNP was extracted, recombination removed using gubbins (v2.4.1)^77^, and a phylogenetic tree was built using fastTree (v2.1.10).^78,79^

### MLST1 *cdtR* SNPs analysis

*cdtR* hits without starting position at the beginning of *C. difficile* contigs were chosen to further examine their nucleotide differences in *cdtR* gene to R20291 and ST1-75. Five such isolates were found either from Patric collection or BioProject PRJEB4556^23,41^, and multiple sequences alignment were performed in Geneious Prime 2022.0.1 with default settings.

### FastANI

Genomic similarity between clinical isolates were calculated using FastANI (v 1.32)^80^ and presented as ANI score.

### Pangenomic analysis of ST1 isolates using Anvi’o

Fifteen circularized genomes of ST1 isolates generated by the Nanopore and Illumina hybrid assemblies were used for pangenomic analysis. Default settings were used based on the Anvi’o workflow for microbial pangenomcis with adjustments for minbit as 0.5 and mcl-inflation as 10.^28,81–83^ Annotations were performed with NCBI Clusters of Orthologous Genes (COG).^84^ Accessory genomes were grouped by gene clusters that are not present in all 15 isolate genomes.

### Prophage identification using PHASTER

Three complete prophages were identified in the genome sequence of strain ST1-75 using PHASTER.^85^ Based on Blastn analyses, the phiCD75-1 prophage corresponds to phi027, a prophage highly conserved among R027 isolates.^86^ The phiCD75-2 prophage is homologous at 99.82% to the well-described phiCD38-2 phage ^29^, whereas the phiCD75-3 prophage seems to be a new phage with no close homolog in public databases. The detection of ORF and gene annotation on phage genomes were performed with PROKKA ^68^, using an E-value threshold of 10E-3 for function assignment. The most recent PHASTER database (last update Dec 22, 2020) was implemented into PROKKA to improve function prediction and overall annotation quality of phage proteins.^85^ The genomes were reorganized so that they started with the terminase gene. Genomic maps were generated using Benchling and finalized with Inkscape v1.2.

### Prophage induction and phage amplification

To confirm the functionality of the phiCD75-2 and phiCD75-3 prophages, induction was performed in TY (2% yeast extract, 3% tryptose, pH 7.4) using two different strategies described previously. The first method was a treatment with 2.5 μg/mL mitomycin C (Novus Biologicals), and the second one was UV irradiation for 10 sec at a wavelength of 365 nm.^87^ The induced cultures were clarified by centrifugation, then filtered on a 0.22 μm filter and the presence of infectious phage particles was confirmed by plating on bacterial lawns of the R20291 epidemic strain using a soft agar overlay method.^87^ Six isolated phage plaques obtained with each induction lysate were picked, diffused in 500 μL phage buffer (50mM Tris-HCl pH 7.5, 100 mM NaCl, 8 mM MgSO_4_) and the identity of the induced phages was determined by PCR using primers specific for phiCD75-2 (LCF0312 5’-AGCGGTATCGGCTTGGTTGTAGAT-3’ and LCF0313 5’-TGCTAGTTTCCTGTCAAGGTCGCT-3’) and for phiCD75-3 (LCF1242 5’-CGACCCACCTAAAGGTATTCA-3’ and LCF1243 5’-GTTCTTTAGTCCAGTTCCCATTTC-3’). Mitomycin C treatment led to induction of the phiCD75-3 prophage whereas UV treatment led to induction of the phiCD75-2 prophage. Each phage was then plaque-purified 3 times to obtain pure phage cultures. Amplification to high titers (> 10^8^ pfu/mL) was done in TY broth using strain R20291 as the host and methods described elsewhere.^29^ Phage genomic DNA was extracted from each lysate and a restriction profile was established using XbaI and HindIII to further confirm the identity of the phages, as done before.^29^

### Creation of new lysogens

New lysogens of strain R20291 carrying either phiCD75-2, phiCD75-3 or both phages were created using a method described before.^29^ Briefly, soft agar overlays containing high titers (> 10^8^ pfu/mL) of phage phiCD75-2, phiCD75-3, or both phages in equivalent amounts were prepared and bacterial dilutions of the wildtype R20291 strain were spread on top of the lawn. Phage-resistant colonies that grew after overnight incubation were picked, re-streaked 3 times on TY agar, and the presence of the respective prophage was detected by PCR and by confirming the phage-resistant phenotype upon re-infection with the corresponding phage(s).

## Quantification and statistical analysis

Results represent means ± SD. Statistical significance was determined by the unpaired t test and one-way ANOVA test. Statistical analyses were performed using Prism GraphPad software v9.3.1 (* p < 0.05; ** p < 0.01; *** p < 0.001; **** p < 0.0001

## Data availability

Whole-genome sequence data were uploaded to National Center for Biotechnology Information (NCBI) Sequence Read Archive (SRA) under BioProject accessions PRJNA885086 and PRJNA595724.

## Acknowledgments

We thank Dr. Craig D Ellermeier for his generously providing us the CRISPR editing system of *C. difficile*, Dr. Olga Soutourina for providing R20291 strain. We thank Melanie Spedale and the animal facilities of the University of Chicago for their help on mouse experiments. We thank Dr. Nhu Nguyen, Dr. Zhenrun Zhang and the rest of the Pamer lab members for helpful discussions. This work was supported by the National Institutes of Health R01 AI095706. The funders had no role in study design, data collection and interpretation, or the decision to submit the work for publication.

## Author contributions

Q. D. and E.G.P conceived the project. Q.D., H.L. and N.D. analyzed the data. Q.D., J.K.S., R. C.S., M.M.A., J.R.G., F.H., R.L.P., V.B., C.M., C.W., A.S. and C. K. performed experiments. M.K. and T.M. isolated *C. difficile* isolates. V.B.Y. and E.S.S. sequenced clinical isolates. Q.D., H.L., L.C.F. and E.G.P interpreted the results and wrote the manuscript.

---

**Supplemental Figure 1.**
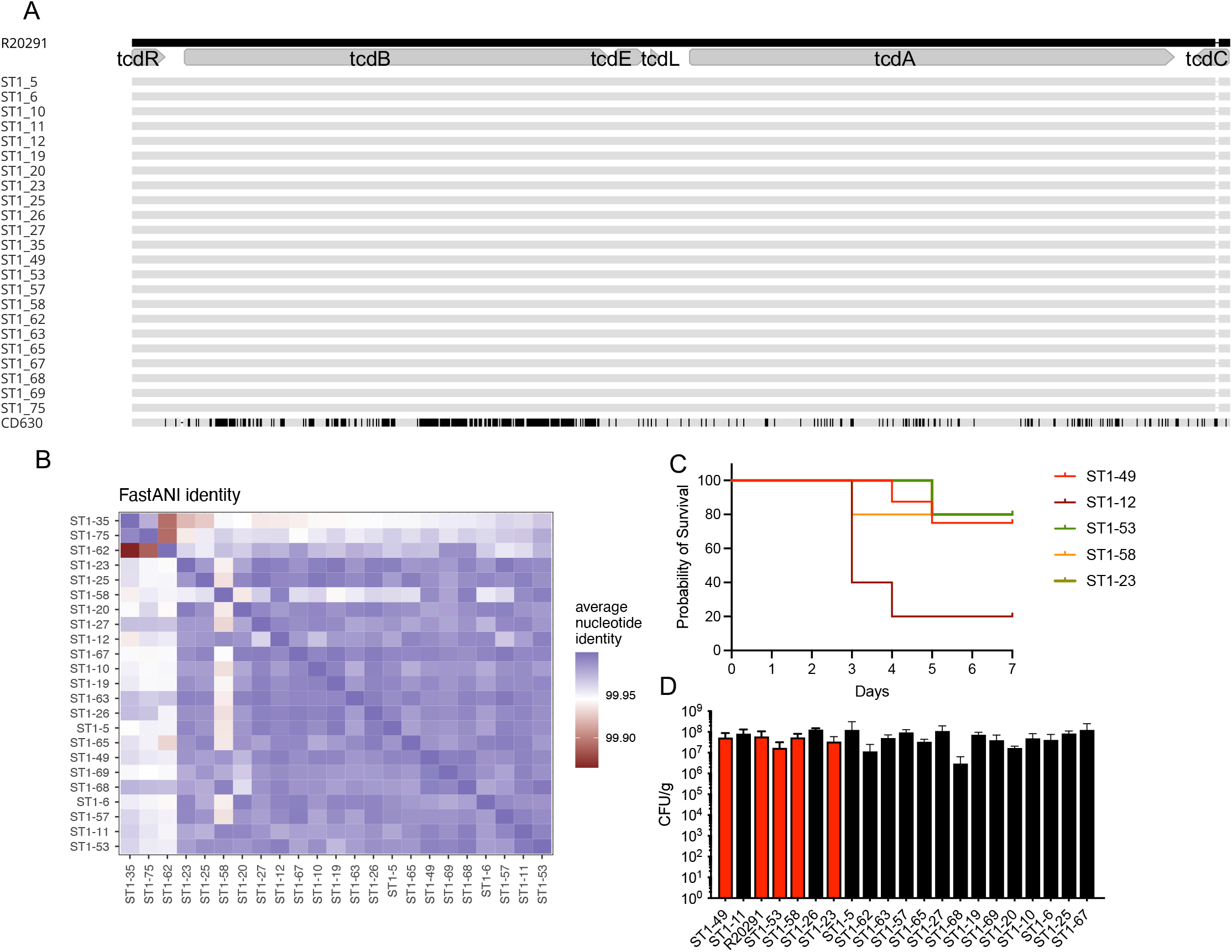
ST1 *C. difficile* strains are closely related. (A) Pathogenicity loci were extracted from whole-genome sequence and multiple sequence alignment was performed for all strains. Each dash indicates one single-nucleotide polymorphism. (B) Average nucleotide identity was calculated with pairs of ST1 isolates. (C) Survival curve of indicated strain over a 7-day time course post infection. (D) Fecal colony-forming units measured by plating on selective agar on 1 day post infection from Figure 1C.

**Supplemental Figure 2.**
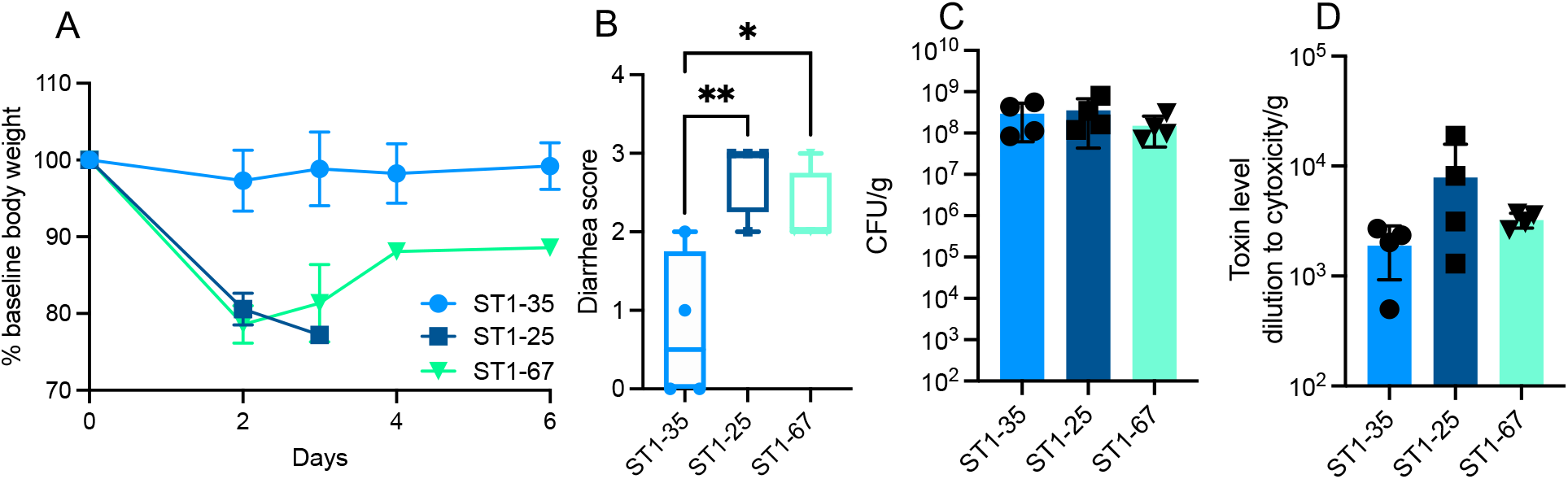
Avirulent *C. difficile* strain demonstrates no virulence in germ-free mice. Germ-free mice (n=4 per group) orally administered with 200 spores of indicated *C. difficile* strains. Daily body weight and acute disease scores were monitored for 6 days post infection. (A) %Weight loss to baseline of mice infected with indicated strains. (B) Diarrhea scores of mice infected with indicated strains on 3 days post infection. (C) Fecal colony-forming units measured by plating on selective agar 1 day post infection. (D) Fecal toxins measured by CHO cell rounding assay 1 day post infection. Statistical significance was calculated by Unpaired t-test, * p < 0.05, ** p < 0.01.

**Supplemental Figure 3.**
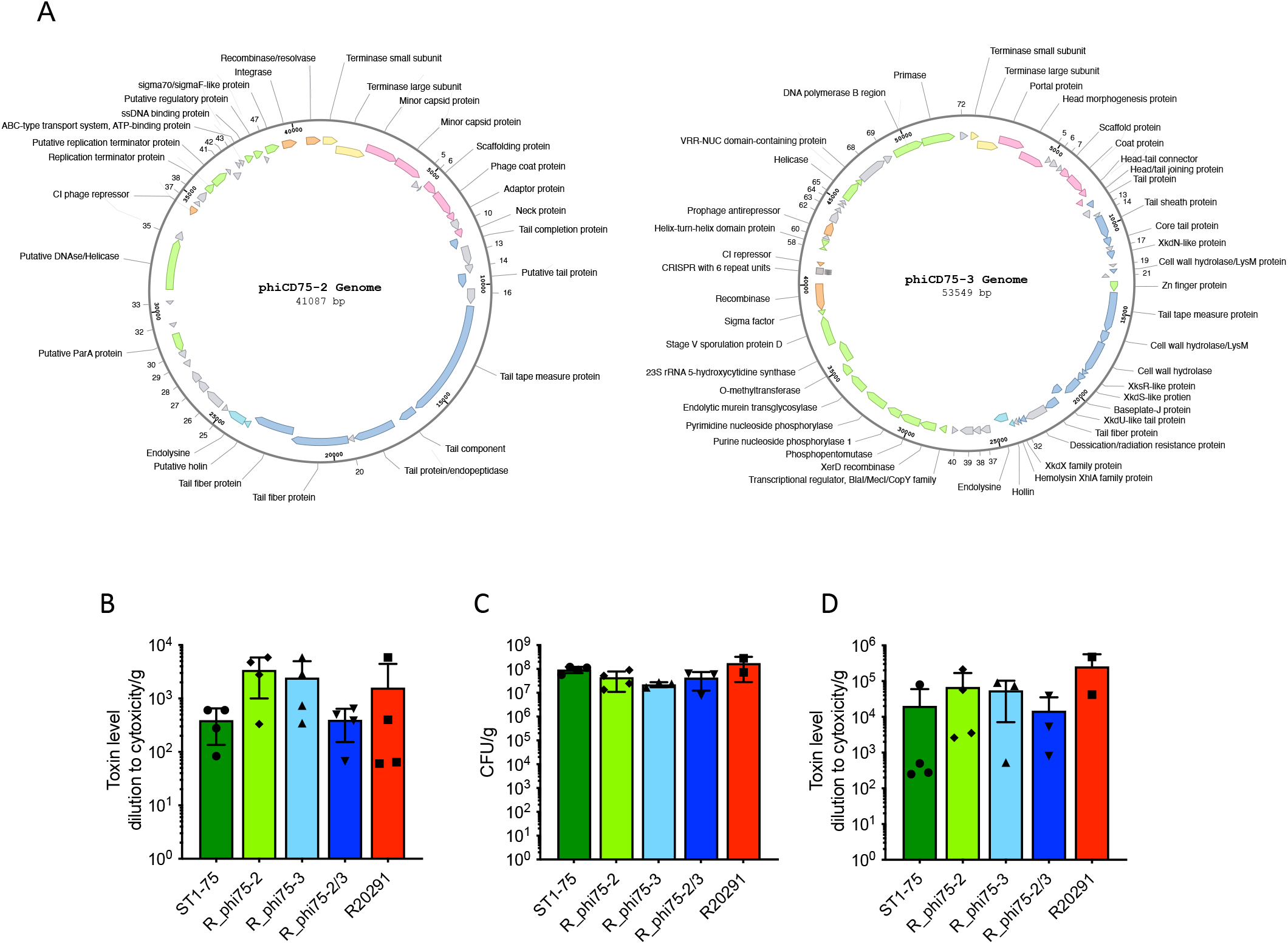
Unique prophages identified in two avirulent *C. difficile* strains. (A) Schematic of prophages identified in avirulent strains. Each arrow represents a coding sequence whose product is indicated, when available. Numbers refer to the CDS when no function could be assigned. Gene products are colored by functional groups; yellow = packaging; red = head morphogenesis proteins; blue = tail morphogenesis proteins; cyan = lysis module; green = gene regulation and DNA replication; orange = lysogeny; grey = other or unknown function. Maps were generated with Benchling and finalized in Inkscape v1.2. (B-D) Wildtype C57BL/6 mice (n=4 per group) were treated with MNV and clindamycin as preciously described. Then, mice were inoculated with 200 *C. difficile* spores via oral gavage. (B) Fecal toxins measured by CHO cell rounding assay 1 day post infection. (C) Fecal colony-forming units measured by plating on selective agar 7 days post infection. (D) Fecal toxins measured by CHO cell rounding assay 7 days post infection.

**Supplemental Figure 4.**
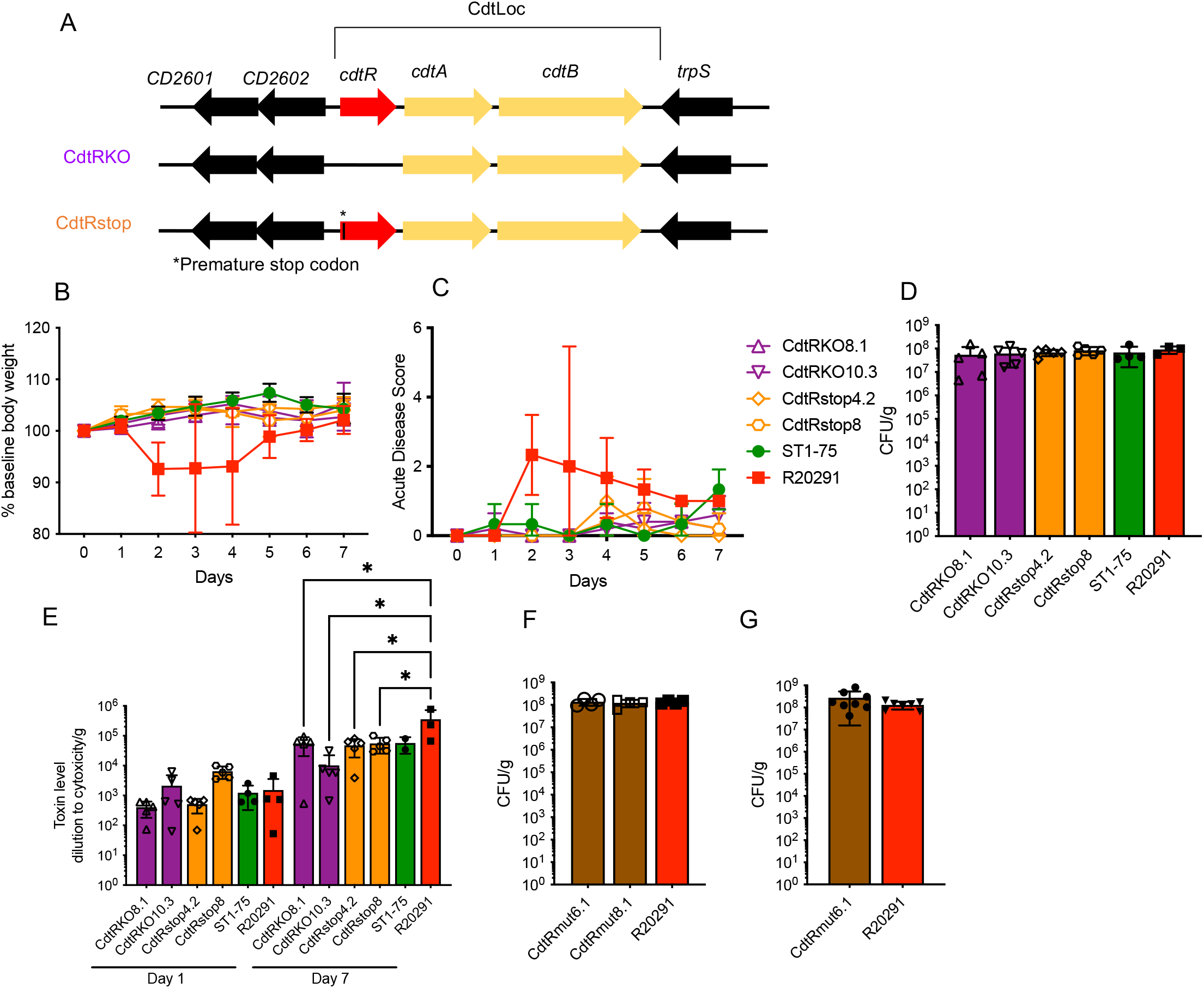
Binary toxin regulator *cdtR* does not impact *C. difficile* colonization in mice. (A) Schematic of *cdtR* mutants generated using R20291 *C. difficile* strain. (B-F) Wildtype C57BL/6 mice (n=3 to 5 per group) were treated with MNV and clindamycin as preciously described. Then, mice were inoculated with 200 *C. difficile* spores via oral gavage. Daily body weight and acute disease scores were monitored for 7 days post infection. (B) %Weight loss to baseline of mice infected with indicated strains. (C) Acute disease scores comprising weight loss, body temperature drop, diarrhea, morbidity of mice infected with indicated strains. (D, F) Fecal colony-forming units measured by plating on selective agar 1 day post infection (E) Fecal toxins measured by CHO cell rounding assay on indicated days. (G) Germ-free mice (n=4) orally administered with 200 spores of indicated *C. difficile* strains. Fecal colony-forming units measured by plating on selective agar 1 day post infection. Statistical significance was calculated by One-way ANOVA, * p <0.05.

**Supplemental Figure 5.**
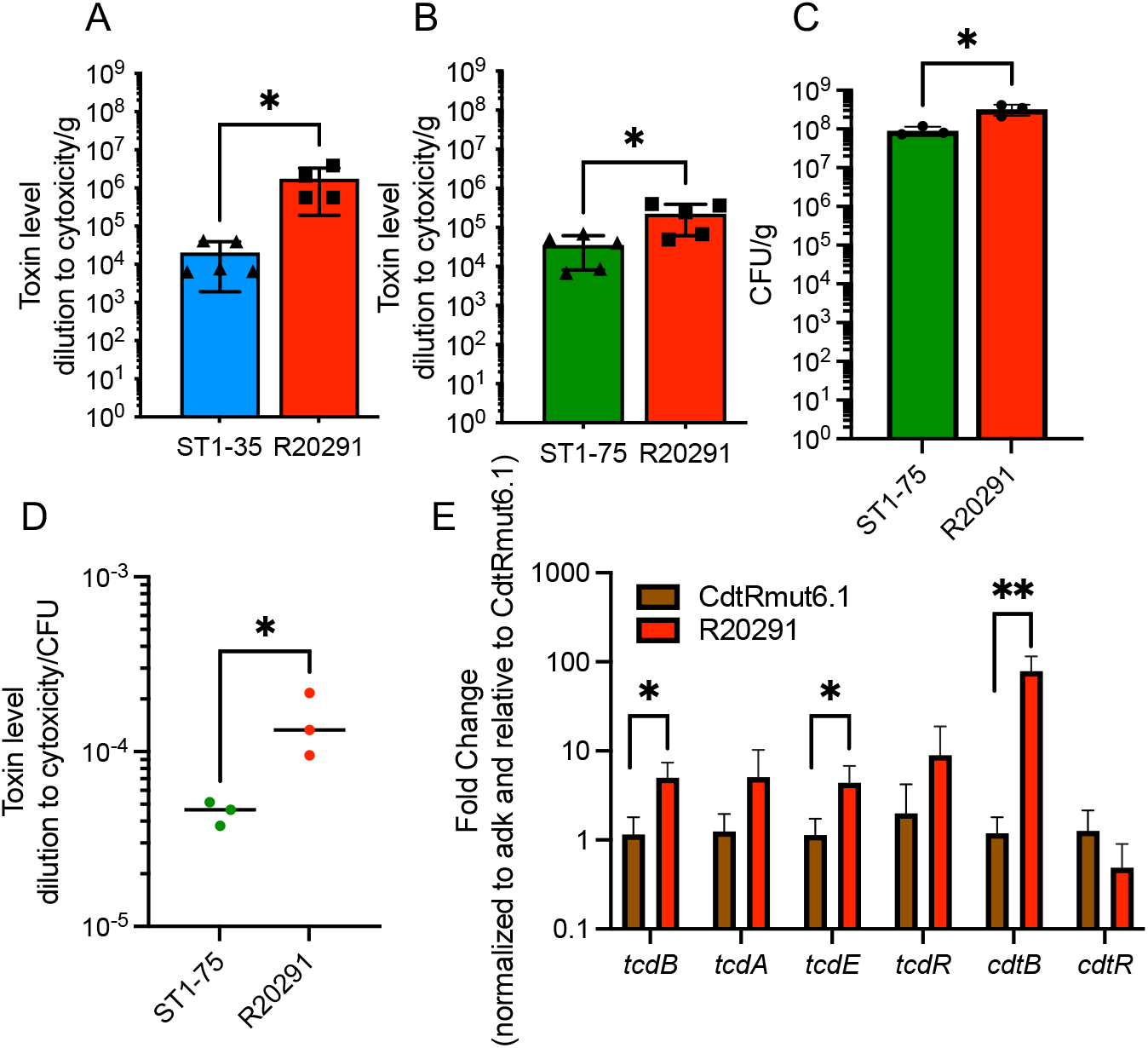
CdtR regulates PaLoc toxins production. (A-B) Wildtype C57BL/6 mice (n=3-5 per group) were treated with MNV and clindamycin as preciously described. Then, mice were inoculated with 200 *C. difficile* spores via oral gavage. (A) Fecal toxins measured by CHO cell rounding assay 7 days post infection for ST1-35. (B) Fecal toxins measured by CHO cell rounding assay 14 days post infection for ST1-75. (C-E) Germ-free mice (n=3-4) orally administered with 200 spores of indicated *C. difficile* strains and cecal contents were harvested at 24 hours post infection. (C) Cecal CFU measured by plating on selective agar. (D) Cecal toxin as in Figure 5A was normalized to CFU. (E) PaLoc and CdtLoc transcripts were measured by RT-qPCR. Transcripts were all normalized to the *adk* and fold change is relative to CdtRmut_6.1 for each of the genes. Statistical significance was calculated by Unpaired t-test, * p < 0.05, ** p < 0.01.

**Supplemental Figure 6.**
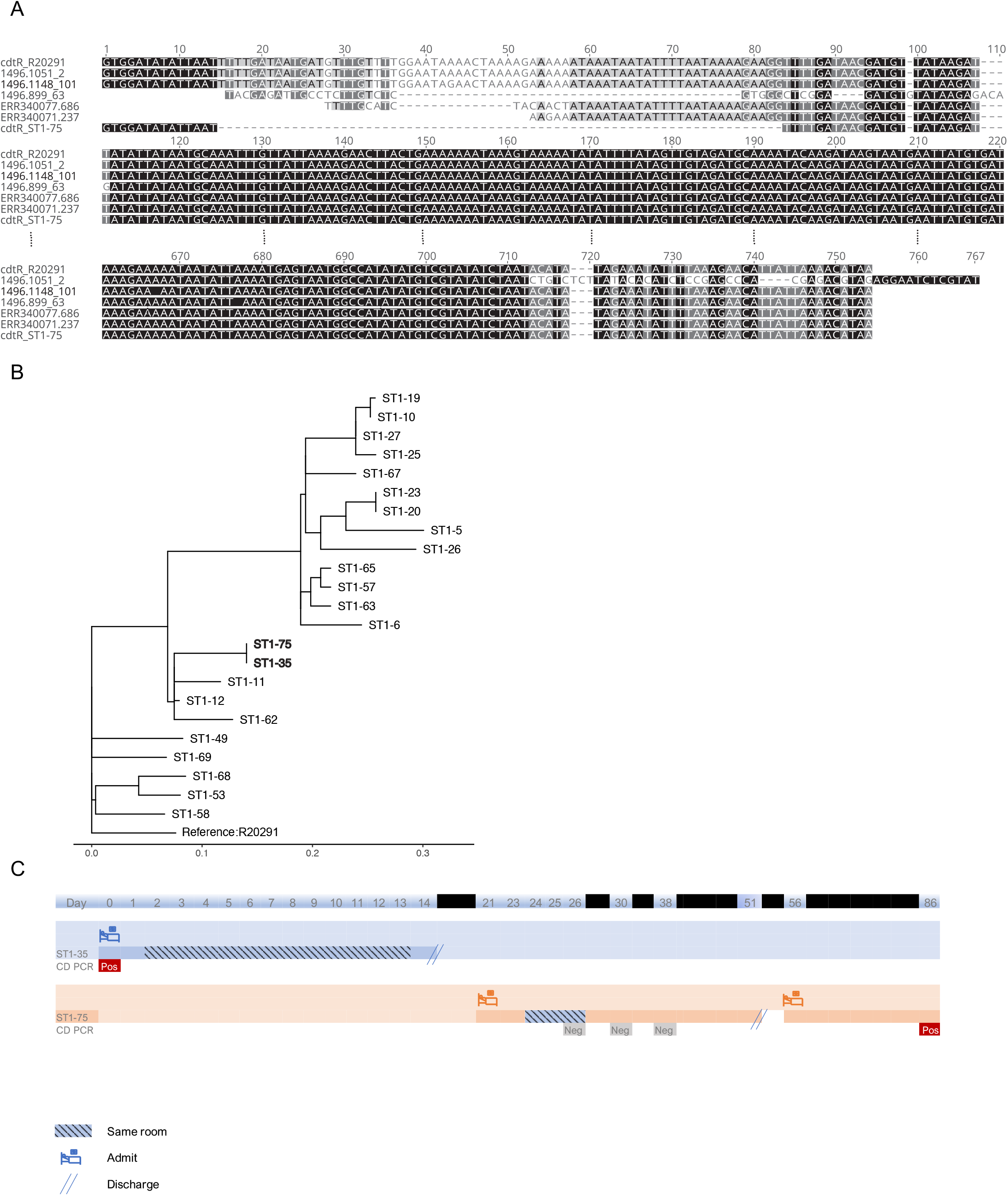
ST1-75/35 harbors unique mutations in *cdtR*. (A) *cdtR* hits without starting position at the beginning of *C. difficile* contigs were chosen to further examine their nucleotide differences in *cdtR* gene to R20291 and ST1-75. Five out of 491 ST1 isolates (from both strain collection databases) were found to have nucleotide variants. (B) Phylogenetic tree build based on core genome snps of ST1 isolates against R20291. (C) Timeline of hospital stay and spatial overlap between the two patients harboring ST1-75 and ST1-35.

